# Transposons contribute to splice-isoform diversity in the *Drosophila* brain

**DOI:** 10.64898/2026.01.22.701052

**Authors:** Malak Choucri, Tina Rainey, Christoph D. Treiber

## Abstract

The extraordinary complexity of the brain depends in part on the vast diversity of mRNA isoforms it expresses, often in a cell-type specific manner. In a recent study, we found that intronic transposable elements (TEs) are spliced into neural transcripts and diversify the splice isoform repertoire of neurons and glia (Treiber and Waddell, 2020). A recent paper by Azad et al. revisits these findings using their TIDAL analysis pipeline applied to our published data (Azad et al., 2024). Their analysis did not find any of the splicing reads we reported, and although they used RT-PCR to test seven of the 264 TE-gene pairs we had previously reported, they failed to validate TE-gene splicing in any of them. Here, we conduct a quantitative analysis of TE exonisation and show that intronic TE insertions are frequently recruited as alternative exons, with exon usage ranging from rare events to near-complete inclusion in transcripts. We implement this analysis in an improved version of our TEChim software, and present clear support for TE-gene splicing at the seven loci tested by Azad et al. We also identify methodological issues in the experimental and computational design of the Azad et al. study that likely explain their failure to detect TE-gene chimeras, while demonstrating that TE-gene splicing can be detected by RT-PCR under appropriate experimental conditions. Together, our data demonstrates that TE splice isoforms are not rare artefacts but measurable and biologically relevant features of the *Drosophila* brain transcriptome that may contribute to the molecular complexity and functional adaptability of the brain.

## Introduction

Neural transcriptomes are shaped by extensive alternative splicing, which generates a large number of mRNA isoforms that drive the structural and functional diversity within the brain. Among the genomic features that can contribute to this process, transposable elements (TEs) are of particular interest. TEs make up a substantial proportion of eukaryotic genomes and provide a rich and diverse reservoir of potential splice sites. However, their contribution to neural transcript diversity remains poorly understood. In our previous work, we showed that many intronic TEs in the *Drosophila* genome are spliced into mature transcripts in the brain, giving rise to chimeric mRNAs that incorporate TE-derived sequence (Treiber and Waddell, 2020). By combining high-coverage genomic DNA (gDNA) sequencing (Treiber and Waddell, 2017) with strand-specific RNA sequencing from the same fly strain, we identified 264 TE-gene pairs with clear, breakpoint-spanning read support for splicing into and out of TEs.

TE insertions are not fixed features of genomes but vary extensively between individuals. In *Drosophila*, TE landscapes differ markedly between laboratory strains and even between individuals of an inbred strain (Treiber and Waddell, 2017). Comparable levels of inter-individual TE polymorphisms are well documented in mammals (Akagi et al., 2008), including humans (Bennett et al., 2004; Ewing and Kazazian, 2011). Importantly, recent large-scale transcriptomic and proteogenomic analyses in humans demonstrate that intronic TE insertions are frequently exonised, giving rise to thousands of recurrent, translated splice isoforms with distinct structural and functional properties (Arribas et al., 2024). Hence, variation in TE landscapes has the capacity to generate inter-individual variation of splice isoform repertoires, and potentially protein structure and abundance. Within the nervous system, TE exonisation therefore provides an intriguing mechanism by which genetic variation in TE landscapes could contribute to inter-individual differences in neural function and behaviour.

Here, we introduce a quantitative analysis of TE exonisation using an extended version of the TEChim pipeline, allowing us to measure the extent to which individual intronic TE insertions are used as alternative exons. We show that TE exonisation efficiency varies widely across loci, and is often substantial, and that these values are reproducible across biological replicates. We then examine why a recent re-analysis of the same data by Azad et al. failed to recover these splice junctions, identifying methodological issues that readily account for their negative results. Finally, we revisit our earlier analysis of TE expression modes to place these splice events within the broader context of autonomous vs. non-autonomous TE expression in the brain.

Together, these lines of evidence support a model in which TE-derived splice isoforms are stable and biologically relevant features of the *Drosophila* brain transcriptome, and, in light of growing evidence from other species, part of a conserved strategy by which eukaryotic genomes diversify gene function.

## Results

### Intronic TE insertions show a broad range of exonisation rates

To quantify how frequently individual TE sequences are used as exons, we extended our TEChim pipeline to calculate, for each breakpoint at an endogenous exon-intron junction, the ratio of TE-derived splice junctions to canonical exon-exon junctions (Figure 1A). For each occurrence, TEChim now reports the relative usage of TE-derived splice junctions immediately downstream (for TE-derived splice acceptor sites) or upstream (for donor sites) of the exon-intron junction compared to canonical exon-exon junctions, providing a simple measure of the TE exonisation efficiency for each event. We applied this analysis to our previously published high-coverage RNA sequencing data (Figure 1B, Supplemental table S1). Many of the TE-gene pairs contain more than one chimeric breakpoint, and we therefore obtained splice ratios for 456 individual TE-gene junctions in total.

**Figure 1.**
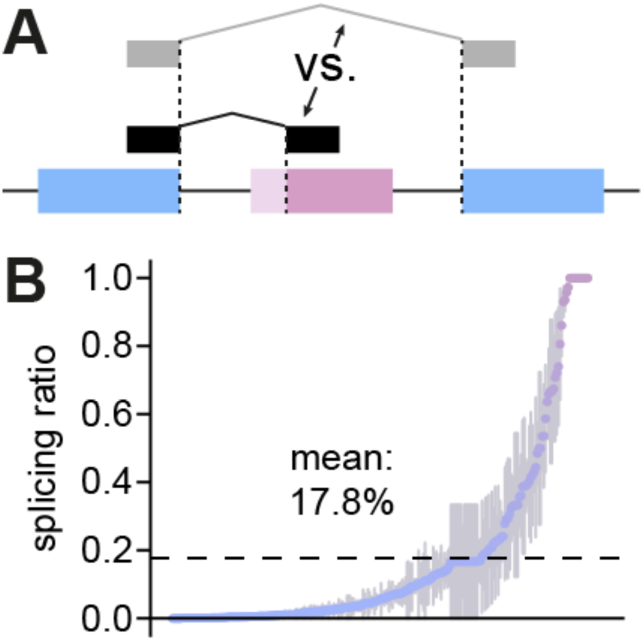
Exonisation rates at intronic TE insertion sites. **(A)** Schematic of the approach used to quantify exonisation rates. For each TE-gene breakpoint at an endogeneous exon-intron junction, TEChim computes the ratio of reads supporting TE-derived splice junctions relative to canonical exon-exon junctions. **(B)** Ranked distribution of TE exonisation efficiencies across intronic and exonic TE insertion sites in the Drosophila brain. For each of the 264 TE-gene pairs, the breakpoint with the highest exonisation ratio is shown. Points represent mean exonisation ratios across six biological replicates. Error bars show SEM. The dashed line indicates the mean exonisation rate across all loci. Junction level exonisation ratios, replicate coverage and SEM values are provided in Supplemental Table S1.

Across these junctions, splice ratios span almost the full dynamic range from the detection floor to 100%. The median junction has a splice ratio of 3%, and a quarter of junctions exceed 15%, indicating that TE-derived exons contribute substantially to the transcript pool at many sites. Where splice ratios can be estimated, values are highly consistent across biological replicates, suggesting that TE exonisation efficiencies are stable properties of individual TE-gene pairs rather than stochastic noise. Overall, these data show that intronic TE insertions in the *Drosophila* brain are frequently recruited as alternative exons, with usage ranging from rare inclusion events to near-constitutive incorporation at the most strongly exonised loci.

### TE-Gene splicing is readily detectable with algorithms that span wider junctions

RNA sequencing is a highly sensitive method to uncover TE-gene chimeric mRNAs (Dobin et al., 2013). Crucially, sequencing reads that span TE-gene breakpoints either exist or they don’t: software such as our TEChim approach, or Azad et al.’s TIDAL does not infer or predict these reads, but simply finds them in the data. In our previous study, TEChim found read support for splicing events into-and out of TEs for 264 TE-gene pairs. For each of these loci we also detected the corresponding TE insertion in genomic DNA (gDNA) from the same fly strain. For TE families represented in multiple TE-gene pairs, the TE-side breakpoints repeatedly mapped to the same internal position within the TE consensus, rather than being scattered along the TE sequence.

One clear example is an intronic antisense insertion of the retrotransposon *roo* inside the *mustard (mtd)* gene, which encodes an ecdysone-regulated puff protein. In abCherry flies, TEChim identified 34 reads upstream and 46 reads downstream of the intron-exon junctions in gDNA, confirming the presence and precise location of the insertion (Figure 2A). In strand-specific RNA-seq data, from six independent biological replicates, TEChim then revealed extensive splicing into-and out of *roo*.

**Figure 2.**
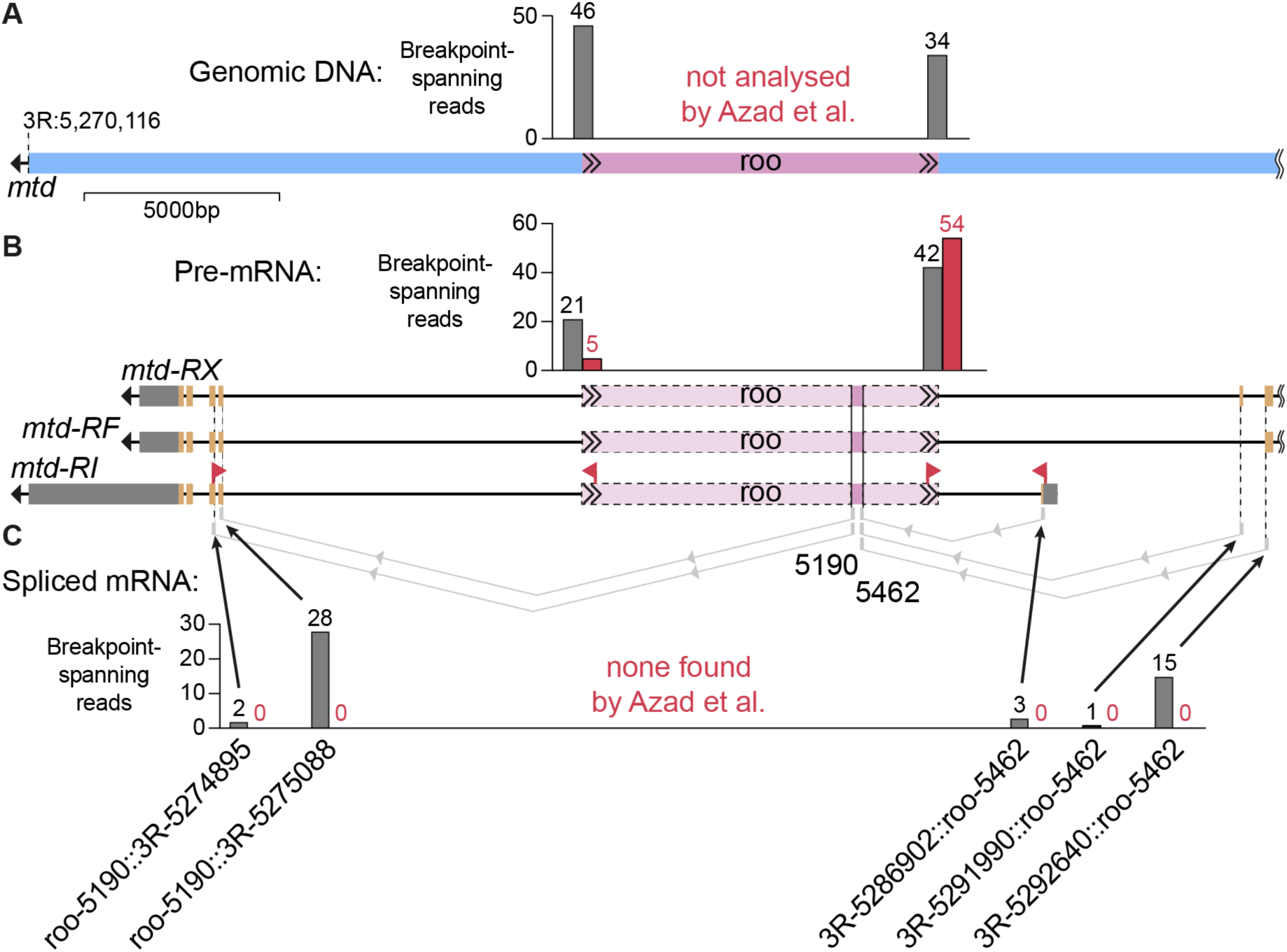
Antisense insertion of roo in mtd introduces cryptic splice donor-and acceptor sites. **(A)** Schematic of the genomic locus of the mustard gene, including a roo insertion present in abCherry flies. Bar graphs show the number of breakpoint-spanning reads identified in gDNA-sequencing data from abCherry flies, using TEChim (grey). **(B, C)** Schematic of mustard transcripts -RX, -RF and -RI, with coding sequences (orange) and UTRs (dark grey). The precise splice acceptor-and donor sites on roo are also shown, and sections of roo that are spliced out are shaded in light magenta. Bar graph in B shows the number of breakpoint-spanning reads found in mRNA from abCherry midbrains by TEChim (grey) and TIDAL (red). Bar graphs in C show the number of reads that support splicing events (indicated by light grey lines) between up-and downstream exons, and two conserved loci within roo (grey bars). Each bar is labelled with the precise location of the breakpoint on roo and the reference genome. TIDAL did not find any of these reads. In addition, red arrows indicate the precise location of RT-PCR primers used by Azad et al. Note that one primer of each primer pair maps onto a section of the TE that is actually spliced out. Figure is drawn to scale and gDNA and mRNA are aligned to each other.

We found 42 reads that spanned the upstream intron-TE breakpoint of *mtd* pre-mRNA in the midbrain and 21 reads that spanned the downstream breakpoint, consistent with unspliced pre-mRNA (Figure 2B). In addition, we detected 49 reads that supported splicing between *mtd* exons and internal positions within *roo*. For example, 15 reads spanned a breakpoint that fell precisely at the end of exon 13 and position 5462 within *roo* (Figure 2C). The antisense sequence leading up to position 5462 resembles the end of an intron and contains a cryptic splice acceptor motif. We also found reads from other upstream exons to the exact same locus within *roo*. Further junctions link position 5190 in *roo* to the next two downstream exons of *mtd*, supported by 28 and 2 reads respectively. Both breakpoints in *mtd* were supported by reads from all six biological replicates. The *roo* sequence contains an imperfect 99bp tandem duplication (two copies, spanning positions 277-475), a 25bp polyA section at 552-576 and a ∼85bp long CAG/AAC microsatellite region at 641-726 (Domínguez, 2021) that could, theoretically, interfere with short read mapping. However, these breakpoints map more than 4kb away from all three regions (Figure 2 – figure supplement 1). We also detected the same breakpoints at 28 other genomic loci (26 for pos. 5190, 23 for pos. 5462; Supplemental table S1). The precise and recurrent patterns of splicing into canonical sequence sections of *roo* across genomic loci and biological replicates make it extremely unlikely that the breakpoint-spanning reads arise from amplification-or mapping artifacts.

Azad et al. applied TIDAL to our RNA-seq data and detected some pre-mRNA breakpoint-spanning reads at the *mtd-roo* locus (Figure 2B, red bars), but they failed to recover any reads spanning splicing events between *roo* and neighbouring exons, and reported similar issues for the other TE-gene pairs they examined (see Figure 2 – figure supplement 2-6). This likely reflects a limitation of their method rather than an absence of splicing. In particular, TIDAL clusters split reads within a narrow window (twice the read length) around candidate junctions, whereas many TE-gene splice junctions fall outside this range. As a result, genuine splice junctions that are readily detected by TEChim fall below TIDAL’s detection criteria. Taken together with our quantitative exonisation analysis, these locus-level examples show that TE-gene splice isoforms are clearly present in the data and are readily recovered when algorithms are configured to capture the relevant splice junctions.

### Strain-specific TE landscapes require genotype-matched validation

TE insertions vary between different lab strains, a well-established phenomenon also documented by the Lau lab (Rahman et al., 2015). In our original study, we used abCherry flies, which were the F1 offspring of MB008b males crossed to UAS-mCherry virgin females. Azad et al. attempted to validate the splice events we found in these flies, but used w1118 and OreR flies, which differ substantially in their genotype. They examined 22 of our loci and found that some insertions were conserved, while others were specific to our abCherry flies. Although this result is not surprising given known differences in TE content between strains, Azad et al. nonetheless interpret it as evidence that “…the majority (∼ 65%) of the predicted TE insertions using RNA-seq as inputs could potentially be false positives”. This interpretation is problematic and potentially misleading. In summary, conclusions about the reproducibility of TE insertions are not valid when based on genotypically mismatched strains.

### Accurate primer design enables robust detection of TE-gene splicing

Strand-specific RNA sequencing enables unbiased, transcriptome-wide detection of splicing events between TEs and genes. Once identified, these events can also be confirmed by a PCR-based approach on cDNA libraries from tissue sample. This requires the careful designing of primers that span both the exon of the gene and the sections of the TE that comprise a given splicing event (Figure 3A).

**Figure 3.**
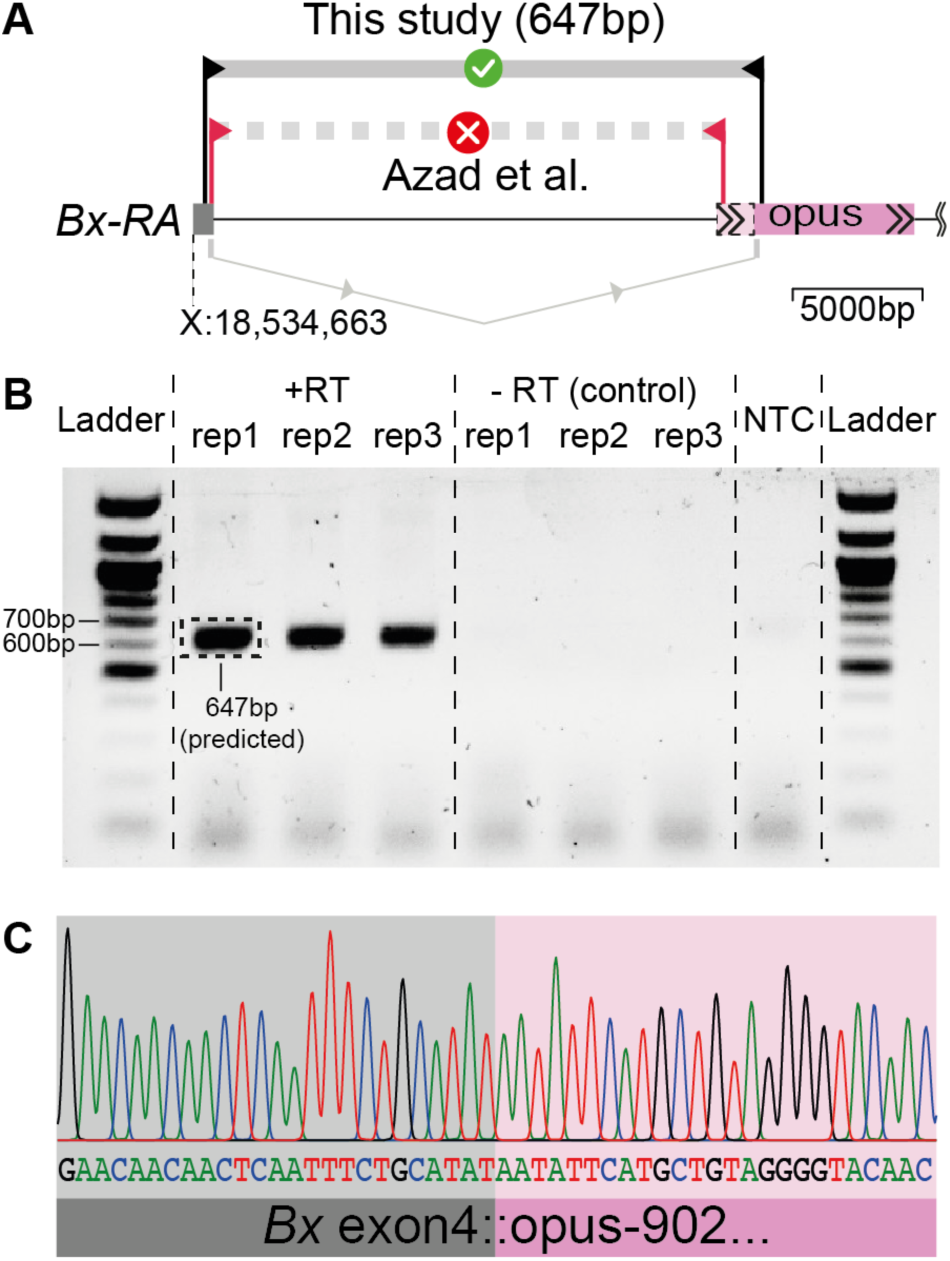
Accurate primer design generates independent breakpoint-spanning fragments that confirm splicing between Beadex and opus. **(A)** Schematic of the genomic locus of Beadex (Bx) - splice isoform -RA, drawn to scale. Primer pairs used by Azad et al. are shown in red (at the precise point where they map). The primers used here are drawn in black (forward primer: CCTGATCTCACGGTCTCTGT; reverse primer: CCTTAATGCCTGTCACCACG). The predicted band size for the spliced transcript is 647bp. **(B)** Electrophoresis gel image of PCR products. The same forward and reverse primers were added to all reactions. The templates for columns 2-4 were three independent biological replicates of +RT samples, columns 5-7 are the matching samples, but their - RT control, and column 8 is the no-template-control (NTC). Columns 1 and 9 contain a Quick-load 100 bp ladder. The gel image is cropped around the edges but otherwise unedited. Dashed box indicates the fragment that was excised and Sanger sequenced. **(C)** Sanger sequencing result of the band from b. Sequence matches the predicted breakpoint between Bx exon 4 and position 902 within opus.

Azad et al. intended to test a total of nine genomic loci with this method. However, they made a number of methodological errors in their designs and therefore failed to validate any of the tested splicing events. For five of the insertions, *mtd/roo* (Figure 2), *Bx/opus* (Figure 3), *CG17698/tabor* (Figure 2 – figure supplement 2), *Pde1C/I-element* (Figure 2 – figure supplement 3), *Dscam2/blood* (Figure 2 – figure supplement 4), primers were placed within the section of the TE that is spliced out. In one case, *Dscam2/blood*, additional primers within the spliced region produced bands that the authors interpret as pre-spliced transcripts. However, the band size in the Azad et al. Fig. 5Diii (bottom right panel) is not consistent with this interpretation: an unspliced amplicon would be 2165bp long, whereas a spliced version, as we reported in our study, should be 616bp. The band appears closer to 616bp than 2165bp, although the authors’ decision to crop and combine gels complicates this interpretation.

Two loci, *Rh7/micropia* and *CaMKII/pogo*, were tested despite the fact we did not report splicing events, and it is therefore unclear which hypothesis was being evaluated. For *mub/opus* (Figure 2 – figure supplement 5), the tested insertion in w1118 flies is located 31kb away from the insertion we reported in abCherry flies and the forward and reverse genomic PCR primers used by Azad et al. flank the wrong site. For *SelR/mdg3* (Figure 2 – figure supplement 6), splicing occurs with splice isoforms *SelR-RG, -RH and -RJ*, but one of the two primer pairs of Azad et al. was specific to a different splice isoform. Their other primer pair is designed correctly. However, the absence of a band highlights the most important issue with their approach. *Mdg3* is located within the 3′UTR of *SelR*, and this primer pair should produce a PCR fragment even in the absence of splicing. The fact that the authors did not detect a band suggests technical problems with their RT-PCR rather than absence of the underlying transcript.

RT-PCR requires careful optimisation, and while the presence of a band (and its confirmation using Sanger sequencing) provides strong evidence for the existence of a certain genomic locus or splicing event, the absence of a band cannot reliably be taken as evidence for its absence (Bustin et al., 2009).

To address the concerns raised by Azad et al. directly, we extracted genomic DNA from Canton-S (CS) flies and designed primer pairs around the upstream-and downstream end of an *opus* insertion within the *Beadex* (*Bx*) gene. We found that CS flies were homozygous for the same insertion that we had found in abCherry flies (Figure 3 – figure supplement 1). Next we extracted RNA from three sets of 40 CS fly brains each, generated cDNA libraries and amplified a fragment consistent with our previous observation, where an upstream exon of *Beadex* is spliced into position 902 of *opus* (Figure 3A, B). We confirmed the band from one of the replicates using Sanger sequencing (Figure 3C), which precisely matched the predicted breakpoint. A minus RT sample and a water control did not produce a band (Figure 3B), confirming the amplified bands did not arise from genomic DNA contamination, or other contaminants in the reagents.

In summary, the combination of genotypically mismatched strains and suboptimal primer design is sufficient to explain Azad et al.’s negative RT-PCR results. Consistent with our quantitative exonisation analysis and locus-level RNA-seq evidence, TE-gene chimeric transcripts are readily detected when genotype and primer placement are matched to the documented TE insertion and splice junction.

### TEs in the brain are predominantly expressed as chimeric transcripts

Our splice-ratio analysis shows that many intronic TE insertions are frequently recruited as alternative exons in neural genes. To place these locus-level observations in a broader context, we next revisit our previous analysis of TE expression modes from our 2020 study (Treiber and Waddell, 2020). In that study, we asked, at the level of TE families, whether TE-derived sequences detected in brain mRNA arise predominantly from autonomous TE expression, or from co-expression with neighbouring genes as chimeric pre-mRNAs and spliced mature mRNAs. Determining the balance between autonomous and non-autonomous TE expression is challenging because TEs are highly repetitive, which complicates the unambiguous assignment of sequencing reads to specific TE copies.

In our 2020 study, we addressed this problem using two complementary approaches. First, we compared the average read depth per nucleotide of individual TE families with the abundance of TE-gene breakpoint-spanning reads for those same families (Figure 4A-C). This analysis indicated that TE expression was most consistent with a non-autonomous origin, arising primarily from co-expression with neighbouring genes. Second, we analysed long terminal repeat (LTR)-containing elements by classifying reads that spanned the LTR and either the internal TE sequence (which can result from both autonomous-and non-autonomous expression), or a flanking genomic region (which can only result from non-autonomous expression). Both approaches supported the conclusion that TEs in the brain are predominantly expressed as part of chimeric mRNAs.

**Figure 4.**
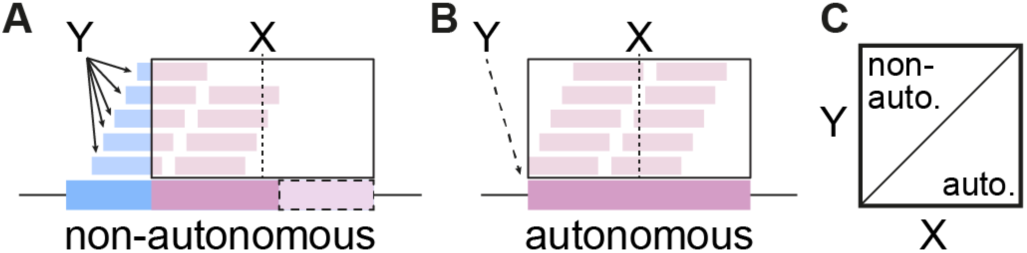
Schematic highlighting method to assess rate of non-autonomous versus autonomous TE expression. **(A,B)** The average read depth per nucleotide within a TE (X) is calculated and compared with the number of breakpoint-spanning reads for the same element (Y). **(C)** Primarily non-autonomously expressed TEs sit in the top left half of the correlation plot, while autonomously expressed TEs are on the bottom right side. The data for this analysis is published in Treiber and Waddell, 2020.

Azad et al. attempted to revisit this question using a different approach. Instead of quantifying the number of reads supporting chimeric vs. non-chimeric transcripts, they compare the rank order of TE families in these two categories. This rank-based comparison lacks a clear biological or statistical interpretation and does not provide a quantitative assessment of TE expression modes. As such, their analysis does not alter the conclusions from our original work that, in the *Drosophila* brain, TE-derived sequences are largely expressed as components of chimeric transcripts, rather than as products of autonomous TE activity.

## Discussion

Here we show that TE-derived splice isoforms are robust and reproducible features of the *Drosophila* brain transcriptome. By quantifying exonisation rates at 456 TE-gene junctions, we find that intronic TE insertions are frequently recruited as alternative exons, with usage ranging from rare events to near-complete incorporation into mature transcripts at a subset of loci. In-depth analyses of candidate TE-gene pairs, together with RT-PCR validation of the *Bx/opus* junction, reinforce the conclusion that chimeric transcripts are readily detectable with carefully-designed experimental and computational approaches. In combination with our previous family-level analysis of TE expression modes, these results support a model in which most TE-derived sequences in the *Drosophila* brain arise from non-autonomous, gene-linked transcripts, rather than widespread autonomous somatic TE expression.

These observations have important implications for how we think about the architecture of neural genes. Intronic TE insertions do not just add passive sequence, but can act as modular exons that are spliced into mature mRNAs (Lev-Maor et al., 2003)Click or tap here to enter text.. Depending on their position within a transcript, TE-derived segments may alter coding sequences, remove protein domains, truncate transcripts, or change untranslated regions that influence RNA stability, localisation or translational control. Because alternative splicing in the nervous system is strongly cell-type specific, TE-driven splice variants provide a straightforward route for generating cell-type specific versions of the same gene product, with distinct TE-derived variants expressed in different neuronal-or glial populations (Brown et al., 2014; Torres-Méndez et al., 2022). Although the present study does not directly address protein output or physiological function, the exonisation rates we report imply that, for many genes, a substantial fraction of transcripts carries TE-derived sequence. This makes it plausible that TE exonisation contributes to the fine-tuning of neural gene function, rather than representing a negligible by-product of noisy splicing.

Beyond cell-type specific regulation, TE exonisation also provides a simple genetic route to variability in neural transcriptomes between individuals and strains. We and others have shown that TE insertion landscapes differ markedly between *Drosophila* strains, and even between individuals. For example, Azad et al. independently confirmed that several insertions we describe are specific to the abCherry background. In a framework where intronic TEs are frequently exonised, such genetic differences translate directly into inter-individual differences in the splice-isoform repertoire of neural genes. One consequence is that genetically distinct flies may express overlapping, but non-identical TE landscapes, which could contribute to variability in neural circuit properties and behaviour between individuals. In the future it will be important to combine TE-aware transcriptomics with single-cell sequencing to determine how TE-derived isoforms are distributed across neural cell types.

A second set of implications concerns methodology. Our re-analysis of the Azad et al. study, especially of their primer sequences, underscores how algorithmic and experimental choices can mask genuine TE-gene splice isoforms. Algorithms like TIDAL, that cluster split reads within narrow genomic windows, or that are not explicitly designed to capture TE-gene architectures, will fail to recover many of the junctions that are visible in the underlying reads. TEChim addresses this by reconstructing full-length fragments from overlapping paired-end reads, mapping these reads to a masked reference genome, and resolving breakpoints with base-pair precision, rather than clustering them within a fixed window. In addition, in TEChim, false discovery rates (FDR) are estimated empirically: expression-matched immobile genetic elements (IGEs) that cannot mobilise in the genome are used to measure the background rate of spurious chimeric reads, and candidate TE-gene breakpoints are only retained once their support exceeds a set FDR. Likewise, RT-PCR validation is only informative when performed in genotype-matched samples using primers that span the reported splice junctions, and when negative results are interpreted in light of assay sensitivity. Emerging direct long-read RNA sequencing technologies offer an attractive complementary approach for validating TE-derived splice isoforms, capturing full-length transcripts and reducing the ambiguity of mapping short reads onto repetitive TE sequence. Such approaches have recently been used to detect analogous TE-gene fusion transcripts in human stem cells and brain tissue (Adami et al., 2025; Garza et al., 2023). Direct RNA sequencing of *Drosophila* brain tissue remains challenging, however, due to the limited RNA input available per sample. Current protocols therefore still require an amplification step, which can itself introduce chimeric artefacts (Treiber and Waddell, 2017). As long-read protocols that require less input material become available, they are likely to further strengthen the characterisation of TE-derived transcript diversity.

More generally, TE transcriptomics requires explicit attention to read mapping, breakpoint calling, strain background and primer design. We suggest that the field would benefit from adopting best-practice guidelines similar to those developed for quantitative PCR (Bustin et al., 2009), with clear standards for reporting TE-aware RNA-seq analyses and breakpoint-level validation.

Several limitations of our study point to concrete next steps. First, our quantitative analysis focuses on a curated set of 264 TE-gene pairs, derived from a single *Drosophila* genotype and bulk midbrain RNA-seq data. Many additional TE insertions likely remain below our detection threshold because of limited read depth or cell-type specific expression. Second, splice ratios in bulk RNA-seq data provide a useful summary of exon usage but do not distinguish between different cell types or developmental stages, and they do not directly reveal whether TE-derived isoforms are translated. Addressing these questions will require TE-aware single-cell sequencing, together with targeted proteomics and genetic manipulations that remove or edit individual TE-derived exons. Finally, while our data show that TE-derived isoforms are reproducible and often abundant, their functional consequences for neuronal physiology and behaviour remain to be tested.

In summary, our work strengthens the view that TEs are not merely genomic parasites or sources of noise, but active contributors to neural gene regulation. Intronic TEs in the *Drosophila* brain are frequently exonised, giving rise to stable chimeric transcripts that expand the splice-isoform repertoire of neuronal and glial genes. By clarifying the methodological requirements for detecting these events and by quantifying their prevalence, we provide a framework for future studies to investigate when and how TE-derived isoforms have been co-opted into neural function.

## Materials and Methods

### TEChim

RNA sequencing data was downloaded from the NCBI BioProject Database (https://www.ncbi.nlm.nih.gov/bioproject/) with the accession number PRJNA588978 (Treiber and Waddell, 2020). Chimeric TE-gene reads were identified using an updated version of TEChim. Paired-end reads were merged with FLASH (Magoč and Salzberg, 2011) into contigs and separated again in-silico to generate new paired-end reads, which were mapped to a repeat-masked reference genome that also contained a single copy of each consensus transposon sequence (and a separate, single section of the long terminal repeat (LTR), where relevant), using STAR (Dobin et al., 2013). Breakpoint-spanning reads were identified using samtools flag filtering (Li et al., 2009), and precise breakpoints were resolved from the corresponding contig using BLAST (Altschul et al., 1990). For mRNA data, candidate breakpoints were then intersected with annotated exon/intron sections and canonical splice sites. The background rate of spurious chimeric reads was estimated using expression-matched immobile genetic elements (IGEs) that cannot mobilise in the genome, and used to define a cutoff for a given False Discovery Rate (FDR). The full pipeline, and detailed documentation are available at https://github.com/charliefornia/TEchim.

### Fly stocks

Wild-type Canton-S flies (Bloomington stock # 64349) were raised on standard brown cornmeal agar food at room temperature and collected at 3-5 days post-eclosion.

### RNA extraction

RNA was extracted using a Promega^TM^ ReliaPrep RNA Tissue Miniprep kit. Midbrains of 20 male and 20 female flies per biological replicate (n = 3 independent biological replicates) were dissected directly into lysis buffer, and tissue was lysed using a Kinematica AG PT1200 Polytron homogenizer. Samples were processed following the standard protocol, including DNAse1 incubation to reduce genomic DNA contamination. Samples were eluted in 22µl nuclease-free water.

### cDNA library preparation

First-strand cDNA synthesis was performed using a Promega^TM^ GoScript Reverse Transcription Mix Oligo(dT) primer. For each biological replicate, the RNA eluate was split into two 10µl aliquots – one was incubated together with GoScript Reverse Transcriptase (“+RT”), and one processed identically, with the same buffers, but without enzyme (“-RT”, the enzyme volume was instead substituted by nuclease-free water).

### Primer design for RT-PCR

Candidate splicing events were first identified using TEChim (see above) and the mRNA and gDNA data were used to reproduce the precise genomic locus, including up-and downstream exons, in ApE (Davis and Jorgensen, 2022). For RT-PCR validation, we designed a gene-specific forward primer upstream of the exon-intron junction in *Bx*, and a TE-specific reverse primer in the region downstream of the cryptic splice acceptor site in *opus* (see Figure 3A), so that the product is only generated across the splice junction, and not from unspliced genomic DNA. Primer design was complicated because *opus*, like most TE sequences, occurs in multiple copies along the *Drosophila* genome. It was therefore necessary to test 2-3 different primer combinations at varying distances from the predicted junction. Every resulting product was Sanger sequenced to confirm it matched the predicted spliced mRNA fragment.

### RT-PCR

cDNA libraries (+RT) and matched -RT controls were analysed in parallel. We followed a standard PCR protocol with APExBT 2X Taq PCR Master Mix, a 1:10 dilution of the stock cDNA libraries, the forward primer: CCTGATCTCACGGTCTCTGT and the reverse primer: CCTTAATGCCTGTCACCACG, with the following conditions: initial denaturation at 94°C for 2 min, followed by 40 cycles of 94°C for 15 s, 56°C for 15 s and 72°C for 30 s, with a final hold at 4°C. In parallel, a no-template control (NTC), with nuclease-free water substituted for cDNA, was included to confirm the absence of contaminating amplification.

### Genomic DNA extraction and PCR

Genomic DNA was extracted from individual Canton-S flies using a rapid single-fly squish preparation. Single flies were homogenised in 50 µl squishing buffer (10 mM Tris-HCl pH 8.0, 1 mM EDTA, 25 mM NaCl) containing 200 µg/ml Proteinase K, added from stock. Homogenates were incubated at 37°C for 30 min, then heat-inactivated at 95°C for 5 min. Debris were pelleted by brief centrifugation, and the supernatant was decanted into a fresh Eppendorf tube. 1 µl of a 1:10 dilution of this supernatant was used directly as template for each gDNA PCR reaction. The opus insertion within the Bx gene was confirmed by a junction spanning PCR on each end. For the upstream end, we used a forward primer in the intron of Bx (GGGTTGCGGCTAAACAGATC) and a reverse primer within the LTR of opus (TGACAACCAACATGCTGTGG), with a predicted fragment size of 888bp if the insertion is present, and no band, if it is not. For the downstream end, we used a forward primer GGATATAAAAGCAGCGGGGC and reverse primer GCGTTTTGTGGCGTGTTAAT, with an expected fragment size of 540bp. PCR produced bands of the expected size in Canton-S, confirmed by Sanger sequencing.

### Sanger-sequencing

PCR products (both from cDNA and gDNA) were loaded on a 1% agarose gel supplemented with SYBRsafe alongside a Quick-Load 100bp DNA Ladder (New England Biolabs), and size separated using standard gel electrophoresis. Bands were excised and purified using a QIAquick Gel Extraction Kit (Qiagen) following the manufacturer’s protocol, with an elution volume of 30µl. Sanger sequencing was performed by Source Bioscience (Nottingham, UK).

## Declaration of generative AI and AI-assisted technologies in the writing process

Generative AI (ChatGPT, OpenAI and Claude, Anthropic) was used during the drafting process to help refine sentence structure, improve clarity, and explore alternative phrasings. All content, data interpretation, and scientific arguments were conceived, written, and verified by the authors.

## Supplemental Figures

**Figure 2 – figure supplement 1:**
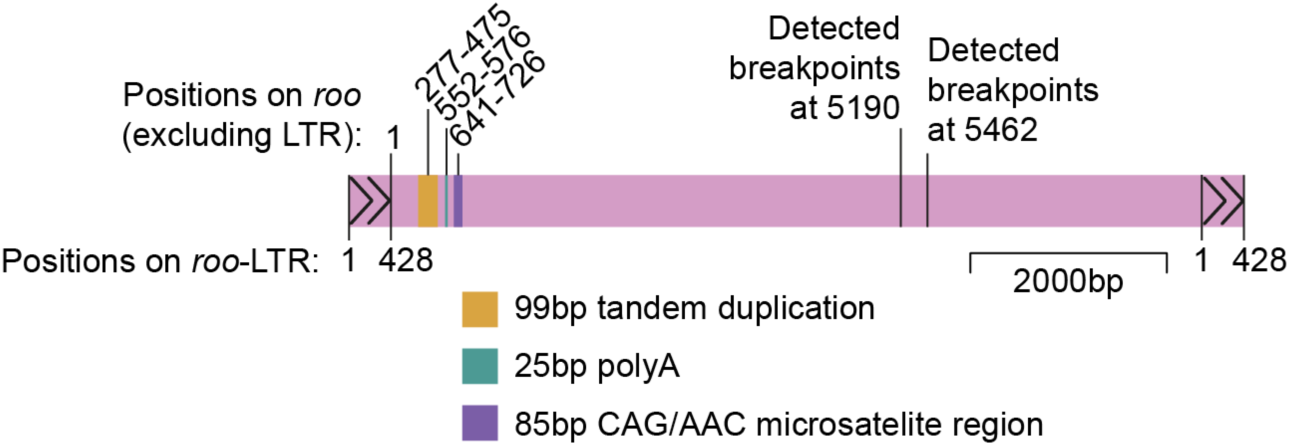
*roo* repeat regions relative to detected breakpoints Schematic of the *roo* consensus sequence, drawn to scale, with positions shown excluding the LTRs (top), and positions of the LTR (bottom). Coloured boxes indicate three annotated repeat features within *roo* (Domínguez, 2021): a 99bp imperfect tandem duplication (positions 277-475), a 25bp poly-A tract (552-576), and an ∼85bp CAG/AAC microsatellite (641-726). The two breakpoints detected in *mtd* (Figure 2) at positions 5190 and 5462, fall more than 4kb downstream of the repeat regions.

**Figure 2 – figure supplement 2:**
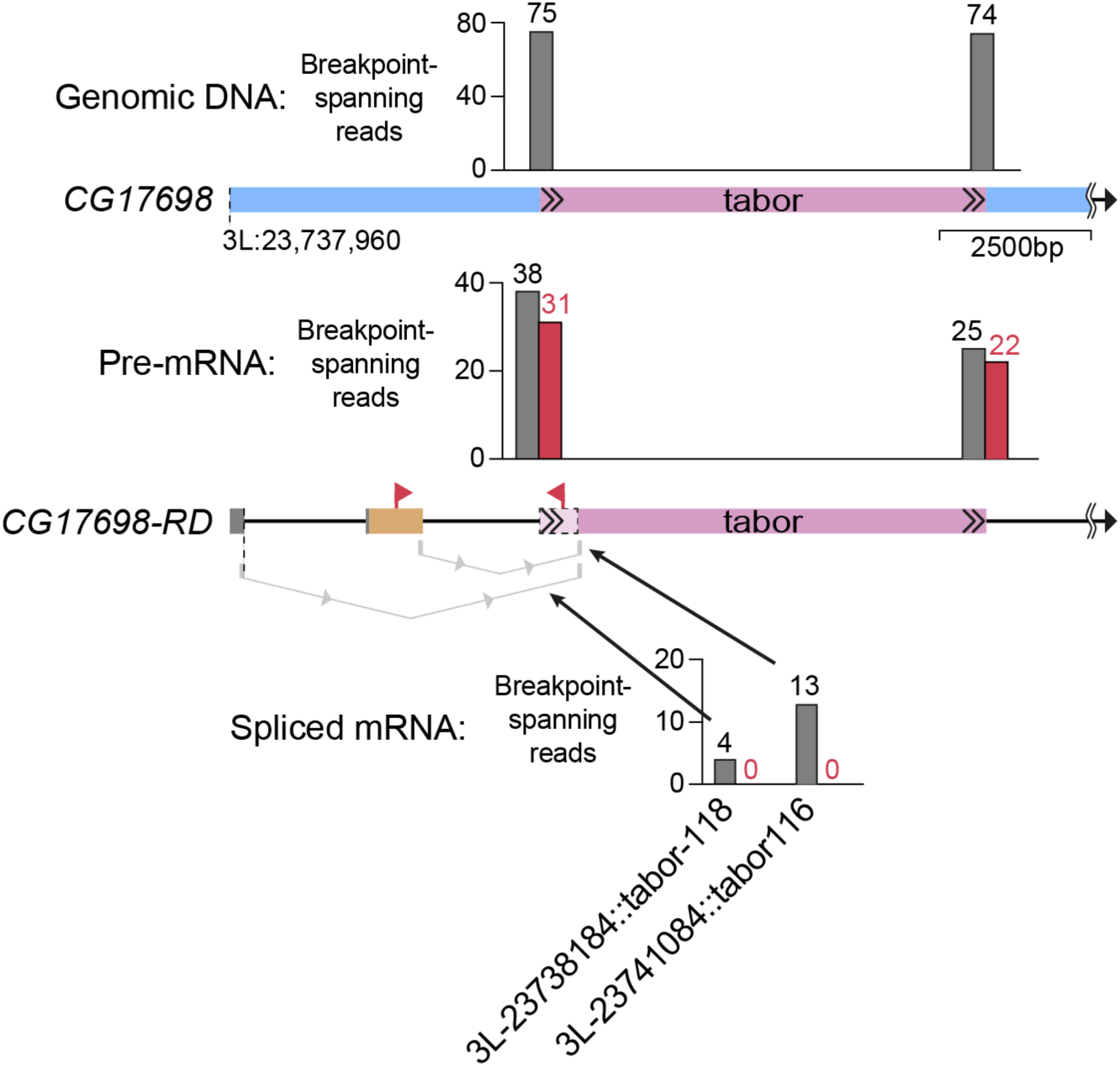
*tabor* insertion in *CG17698* Upper panel shows a schematic of the genomic locus of the *CG17698* gene, including a *tabor* insertion present in abCherry flies. Bar graphs at the top show the number of breakpoint-spanning reads identified in gDNA-sequencing data from abCherry flies, using TEChim. Lower panel shows a schematic of the *CG17698-RD* transcript, with coding sequences (orange) and UTRs (dark grey). The splice acceptor site on *tabor* is also shown, and the section of *tabor* that is spliced out is shaded in light magenta. Bar graphs show the number of breakpoint-spanning reads found in mRNA from abCherry midbrains using TEChim (grey) and TIDAL (red). Bar graphs at the bottom show the number of reads that support splicing events (indicated by light grey lines) between two downstream exons and a conserved locus within *tabor* (grey bars). Each bar is labelled with the precise location of the breakpoint on *tabor* and the reference genome. TIDAL did not find any of these reads. In addition, red arrows show precise location of RT-PCR primers used by Azad et al. Note that one primer maps onto a section of the TE that is actually spliced out. Figure is drawn to scale and gDNA and mRNA are aligned to each other.

**Figure 2 – figure supplement 3:**
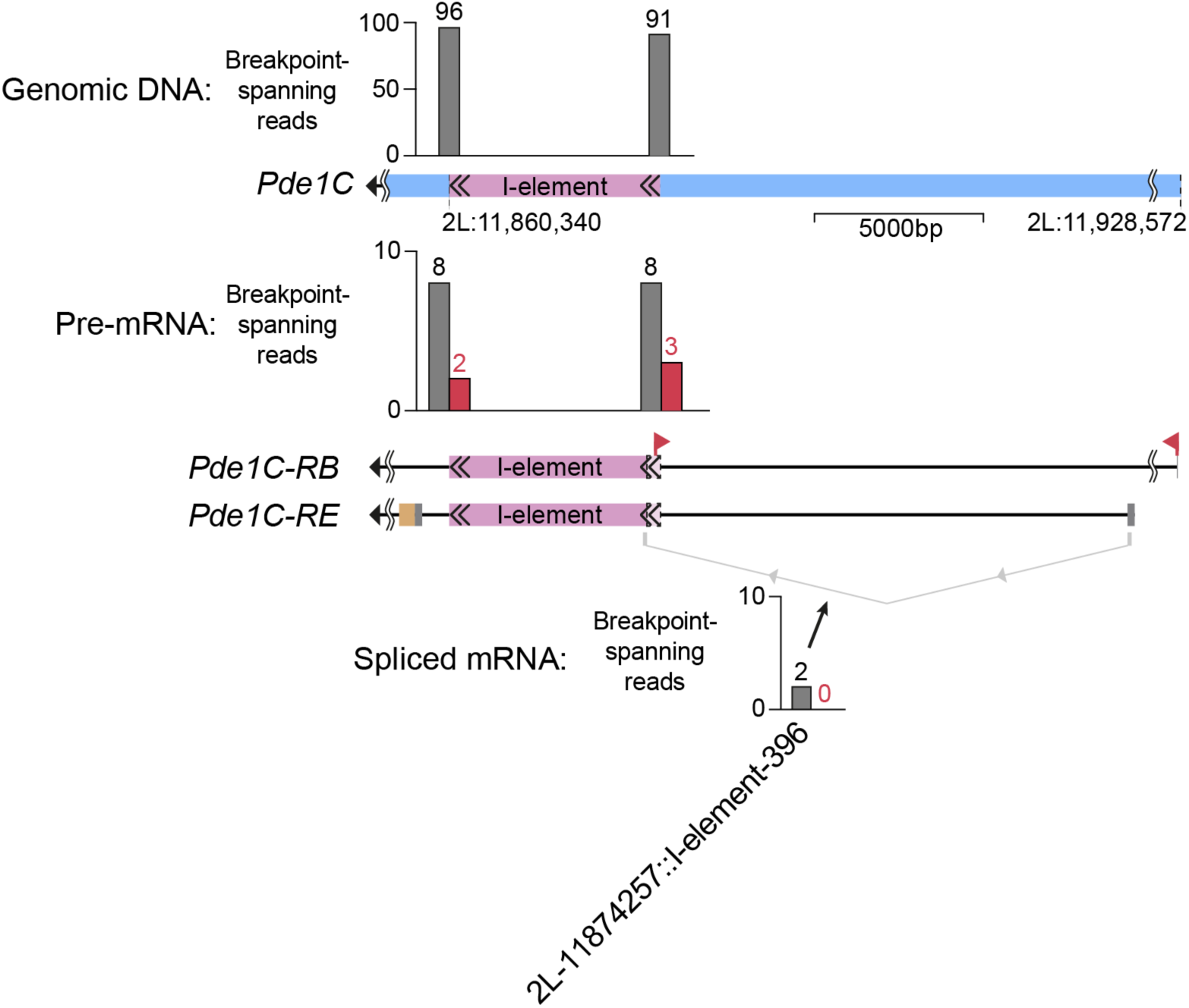
*I-element* insertion in *Pde1C* Upper panel shows a schematic of the genomic locus of the *Pde1C* gene, including an *I-element* insertion present in abCherry flies. Bar graphs at the top show the number of breakpoint-spanning reads identified in gDNA-sequencing data from abCherry flies, using TEChim. Lower panel shows a schematic of the Pde1C-RB and -RD transcripts, with coding sequences (orange) and UTRs (dark grey). The splice acceptor site on the *I-element* is also shown, and the section of the *I-element* that is spliced out is shaded in light magenta. Bar graphs show the number of breakpoint-spanning reads found in mRNA from abCherry midbrains using TEChim (grey) and TIDAL (red). Bar graph at the bottom shows the number of reads that support splicing events (indicated by light grey line) between the upstream exon and a conserved locus within the *I*-element (grey bar). The bar is labelled with the precise location of the breakpoint on the *I-element* and the reference genome. TIDAL did not find any of these reads. In addition, red arrows show precise location of RT-PCR primers used by Azad et al. Note that one primer maps onto a section of the TE that is actually spliced out and the other primer to an exon of another splice isoform than reported in abCherry flies. Figure is drawn to scale and gDNA and mRNA are aligned to each other.

**Figure 2 – figure supplement 4:**
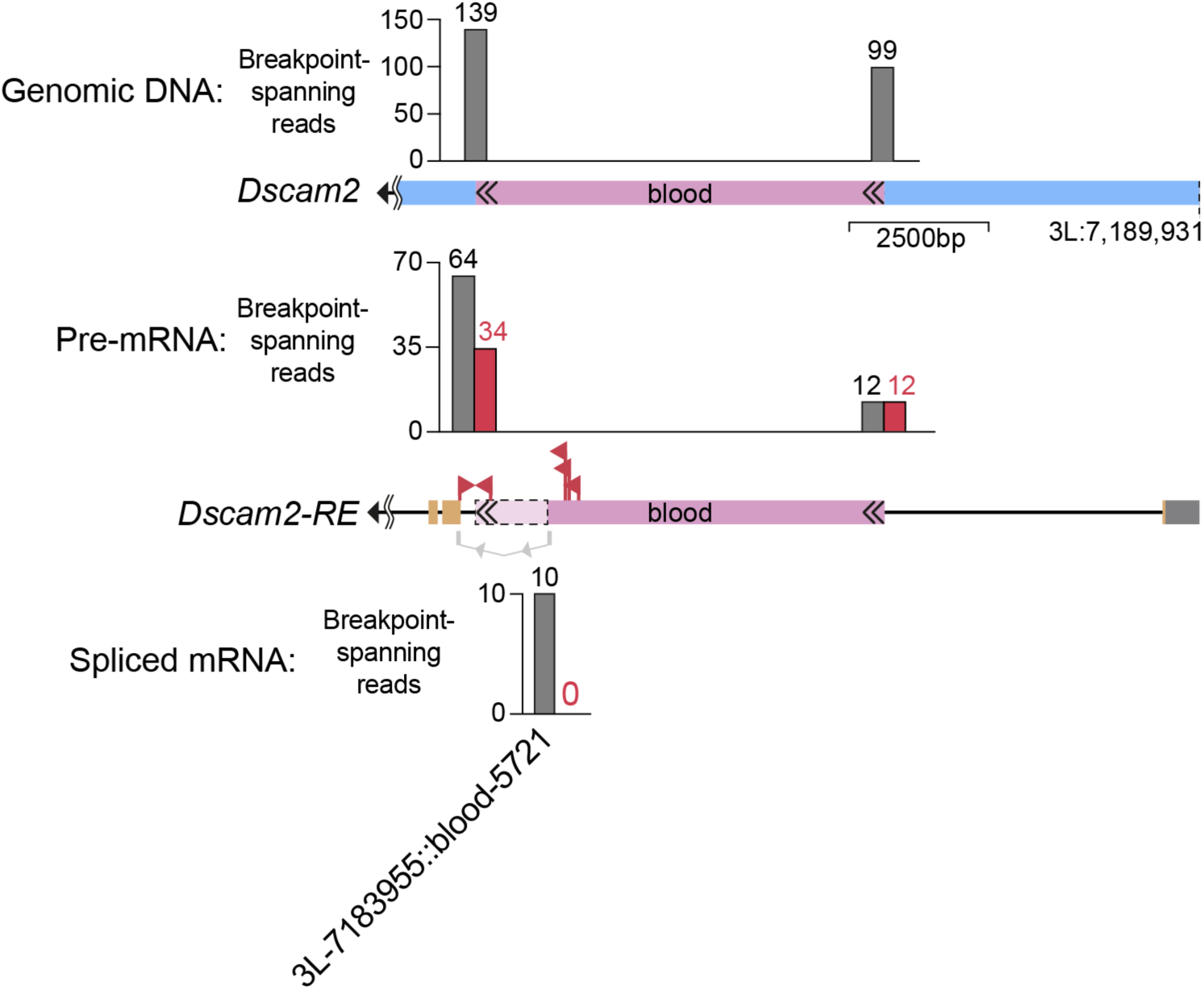
*blood* insertion in *Dscam2* Upper panel shows a schematic of the genomic locus of the *Dscam2* gene, including a *blood* insertion present in abCherry flies. Bar graphs at the top show the number of breakpoint-spanning reads identified in gDNA-sequencing data from abCherry flies, using TEChim. Lower panel shows a schematic of the *Dscam2-RE* transcript, with coding sequences (orange) and UTRs (dark grey). The splice donor site on *blood* is also shown, and the section of *blood* that is spliced out is shaded in light magenta. Bar graphs show the number of breakpoint-spanning reads found in mRNA from abCherry midbrains using TEChim (grey) and TIDAL (red). Bar graph at the bottom shows the number of reads that support splicing events (indicated by light grey line) between the downstream exons and a conserved locus within *blood* (grey bar). The bar is labelled with the precise location of the breakpoint on *blood* and the reference genome. TIDAL did not find any of these reads. In addition, red arrows show precise location of RT-PCR primers used by Azad et al. Note that one primer maps onto a section of the TE that is actually spliced out. Figure is drawn to scale and gDNA and mRNA are aligned to each other.

**Figure 2 – figure supplement 5:**
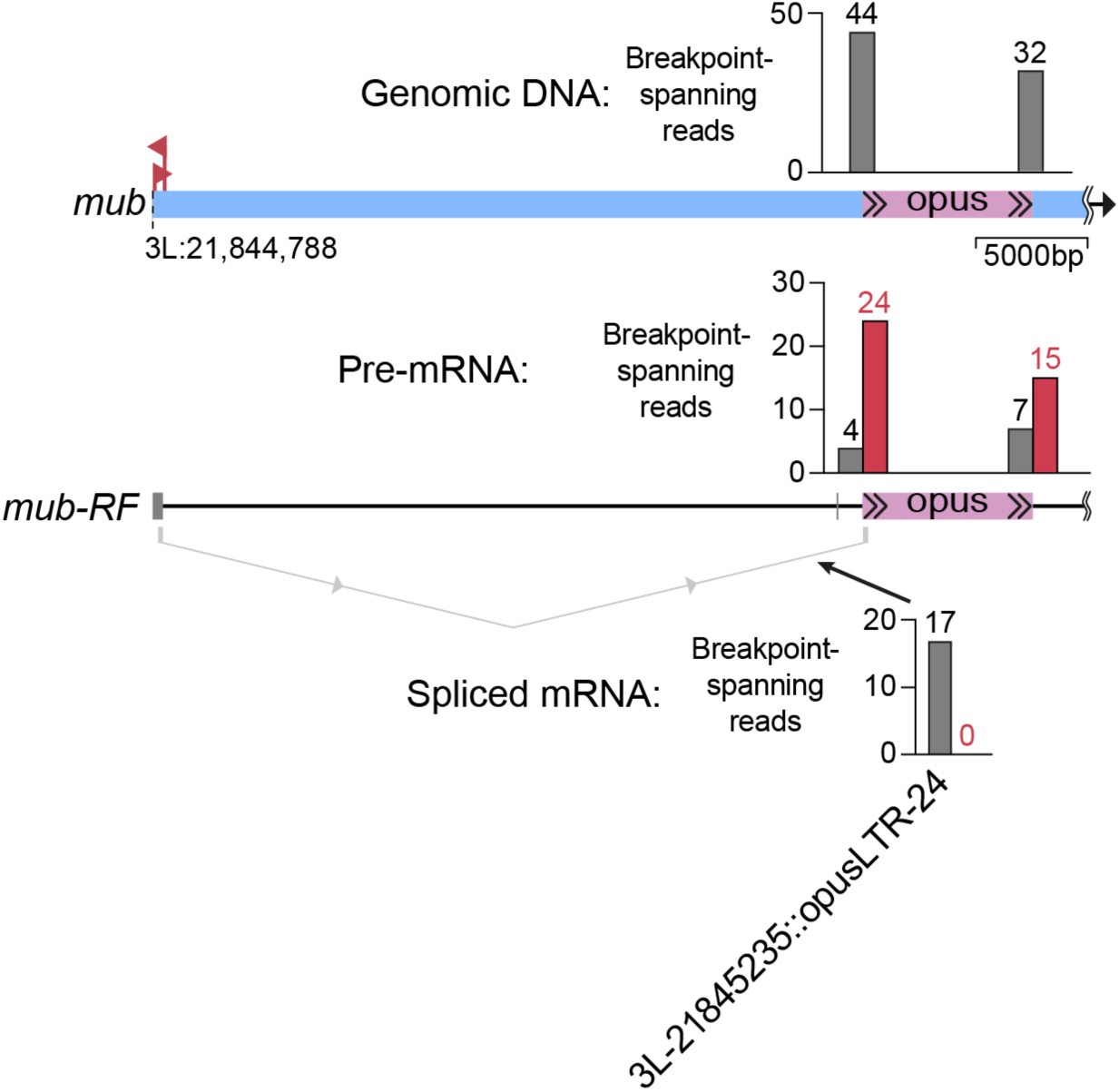
*opus* insertion in *mub* Upper panel shows a schematic of the genomic locus of the *mub* gene, including an *opus* insertion present in abCherry flies. Bar graphs at the top show the number of breakpoint-spanning reads identified in gDNA-sequencing data from abCherry flies, using TEChim. Lower panel shows a schematic of the *mub-RF* transcript, with UTRs (dark grey). The splice donor site on *opus* is also shown. The first 24 nucleotides of *opus* are spliced out, but this cannot be visualised at the current scale. Bar graphs show the number of breakpoint-spanning reads found in mRNA from abCherry midbrains using TEChim (grey) and TIDAL (red). Bar graph at the bottom shows the number of reads that support splicing events (indicated by light grey line) between the downstream exons and a conserved locus within *opus* (grey bar). The bar is labelled with the precise location of the breakpoint on *opus* and the reference genome. TIDAL did not find any of these reads. In addition, red arrows show precise location of PCR primers used by Azad et al. to genotype their w1118 flies. Note that these primers flank a region ∼31kb upstream of the *opus* insertion in abCherry flies. Figure is drawn to scale and gDNA and mRNA are aligned to each other.

**Figure 2 – figure supplement 6:**
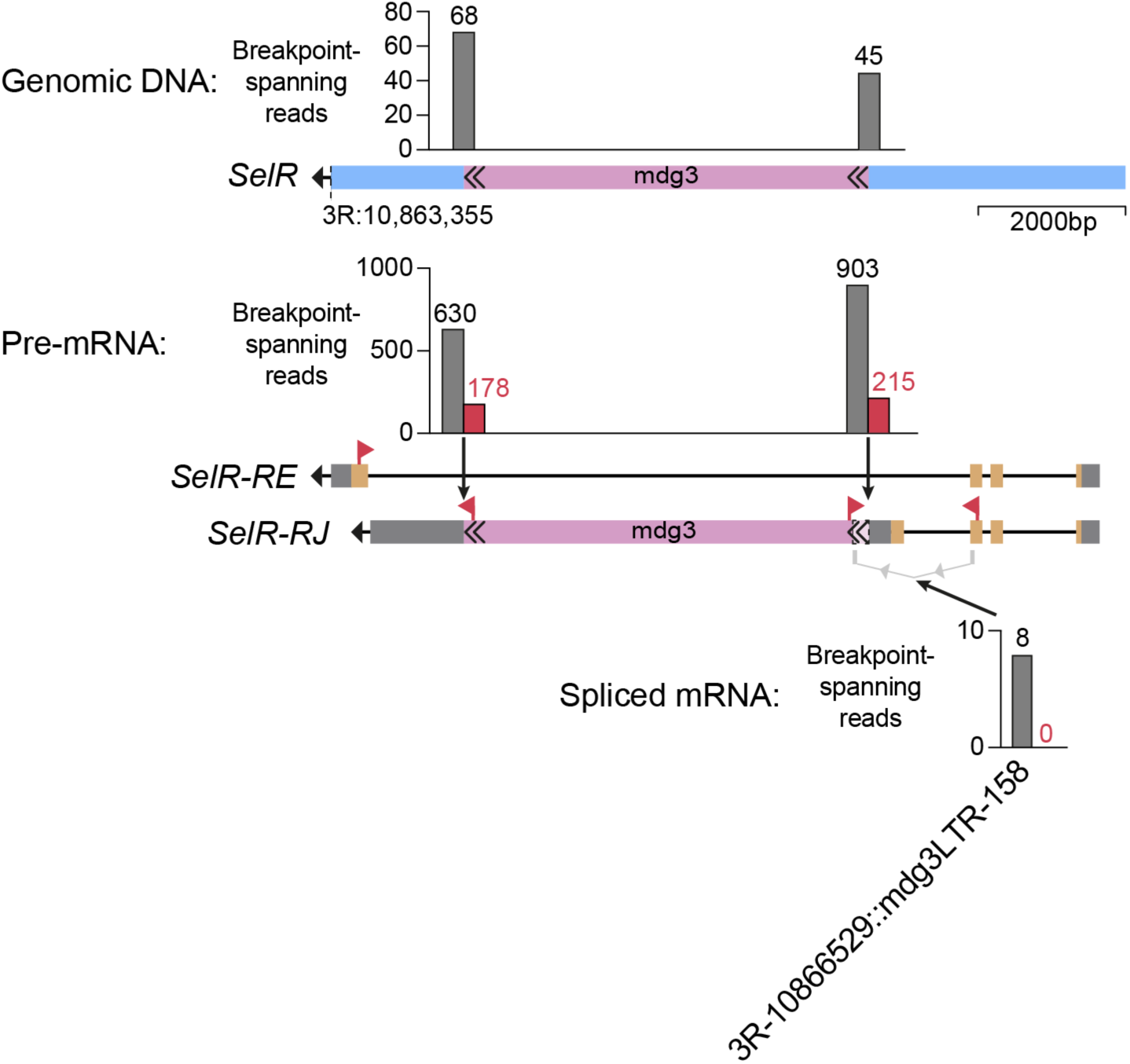
*mdg3* insertion in *SelR* Upper panel shows a schematic of the genomic locus of the *SelR* gene, including an *mdg3* insertion present in abCherry flies. Bar graphs at the top show the number of breakpoint-spanning reads identified in gDNA-sequencing data from abCherry flies, using TEChim. Lower panel shows a schematic of the *SelR-RE and -RJ* transcripts, with coding sequences (orange) and UTRs (dark grey). The splice acceptor site on *mdg3* is also shown, and the section of *mdg3* that is spliced out is shaded in light magenta. Bar graphs show the number of breakpoint-spanning reads found in mRNA from abCherry midbrains using TEChim (grey) and TIDAL (red). Bar graph at the bottom shows the number of reads that support splicing events (indicated by light grey line) between the upstream exon and a conserved locus within *mdg3* (grey bar). The bar is labelled with the precise location of the breakpoint on *mdg3* and the reference genome. TIDAL did not find any of these reads. In addition, red arrows show precise location of RT-PCR primers used by Azad et al. Note that one primer maps onto a section of another splice isoform then reported in abCherry flies. Figure is drawn to scale and gDNA and mRNA are aligned to each other.

**Figure 3 – figure supplement 1:**
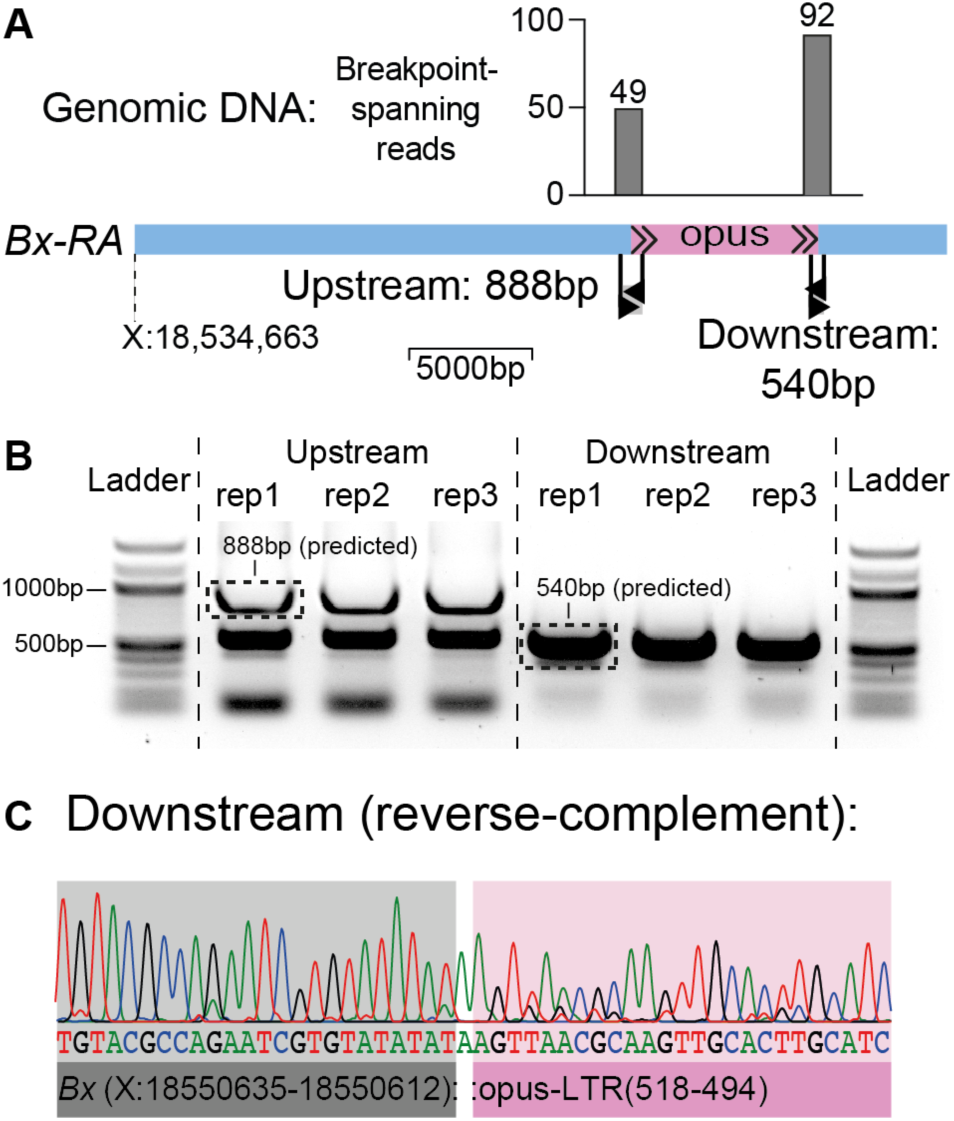
Targeted PCR and Sanger sequencing confirms the *opus* insertion in *Bx*, previously identified by whole-genome sequencing of abCherry flies (A) Schematic of the genomic locus of *Bx*-splice isoform -RA, drawn to scale. Bar graphs at the top show the number of breakpoint-spanning reads identified in gDNA-sequencing data from abCherry flies, using TEChim. Primer loci are shown in black. The predicted fragment size for a genomic locus with the *opus* insertion is 888bp for the upstream and 540bp for the downstream breakpoint. (B) Electrophoresis gel image of PCR products. The templates for columns 2-4 and 5-7 are three independent biological replicates of genomic DNA extracted from single CS flies. Columns 1 and 8 contain a Quick-load 100 bp ladder. The gel image is cropped around the edges but otherwise unedited. Dashed boxes indicate the fragments that were excised and Sanger sequenced. (C) Sanger sequencing result of the downstream band from B. The sequence matches the predicted breakpoint between *Bx* and the end of the opus LTR. Note that the Sanger sequence was done with the reverse primer, hence the sequence shown is the reverse complement to the positive strand of *Bx* and *opus*. Sanger sequencing of the upstream 888bp band (also indicated in B) confirmed flanking *Bx* and *opus* sequence, but did not resolve a continuous read across the breakpoint, likely due to a tandem repeat near the *opus* LTR boundary (see published response to reviewers for more details).

